# Association between genetic clades and cancer prevalence suggested by French-wide study of oncogenic small ruminant beta-retroviruses diversity

**DOI:** 10.1101/2024.07.18.604097

**Authors:** Riocreux-Verney Benjamin, Verneret Marie, Diesler Rémi, Dolmazon Christine, Gineys Barbara, Cadoré Jean-Luc, Turpin Jocelyn, Leroux Caroline

## Abstract

ENTV (Enzootic Nasal Tumor Virus) and JSRV (Jaagsiekte Sheep Retrovirus) are β-retroviruses responsible for respiratory cancers in sheep and goats. In this study, we analyzed the genetic features of the sheep and goat β-retrovirus (29 JSRV and 24 ENTV strains) circulating in France to identify molecular signatures associated with disease severity in flocks. We developed a highly specific PCR to amplify and sequence exogenous targeted regions or near full length proviruses based on limited discriminating motifs along their genomes. The phylogenetic reconstructions based on the LTR and *env* regions suggest that one major strain is circulating on the French territory for ENTV-1 and ENTV-2 while not clustering with already published Spanish, Canadian or Chinese strains. JSRV strains circulating in French sheep flocks were distributed in 2 distinct genetic clades clustering with sequences originating from North America, Africa and United-Kingdom. JSRV clade I was found to be associated with a higher incidence of cancer in French flocks. Specific motifs spanning the entire JSRV genome particularly in the LTRs and in the intracytoplasmic domain of the envelope were detected between the two genetic subtypes.

This work represents the first nationwide study describing the circulation of the three closely related β-oncogenic retrovirus JSRV, ENTV-1 and ENTV-2 in French sheep and goat flocks. Better characterization of strain genetics is a critical step in monitoring circulating β retrovirus, especially those associated with higher cancer incidence in small ruminants.

## 1 Introduction

JSRV (Jaagsiekte Sheep Retrovirus) and ENTV (Enzootic Nasal Tumor Virus) are two oncogenic β-retroviruses responsible for respiratory tumors of the lung and of the nasal mucosa respectively in domestic small ruminants. These diseases have been reported in many regions of the world, including Western Europe, Africa, Asia and North America (York et al., 1992; Bai et al., 1999; DeMartini et al., 2001; Ortín et al., 2003; Caporale et al., 2005; Walsh et al., 2010; Lee et al., 2017; Sid et al., 2018; Nor El Houda et al., 2020; Mao et al., 2022). JSRV-induced cancer is associated with persistent cough, dyspnea, cachexia and fluid production by the tumoral epithelial cells. Tumors arise from the transformed alveolar and bronchiolar epithelial cells. ENTV type 1 in sheep and type 2 in goats induce tumors by uncontrolled proliferation of nasal epithelial cells, resulting to respiratory distress and to nasal discharge, which may be associated with facial deformities and osteolysis. These respiratory cancers lead to rapid death (within a few weeks) after the onset of the first clinical signs; thus, resulting in significant economic losses and a major impact on animal welfare. These viruses, are transmitted by air, contact, *via* colostrum and milk (Grego et al., 2008) and *in utero* (Caporale et al., 2005).

The envelope (Env) of JSRV and ENTV has been shown to be the major oncogenic determinant. Env is cleaved into the surface (SU) and transmembrane (TM) glycoproteins, the former interacting with the cellular receptor Hyal2 (hyaluronidase 2) for viral entry and the latter responsible for cell transformation *in vitro* and *in vivo* (Palmarini et al., 2001; Alberti et al., 2002; Caporale et al., 2006; Wootton et al., 2006; Cousens et al., 2007; Maeda et al., 2021). Through its intracytoplasmic tail, JSRV and ENTV Env deregulates cell proliferation pathways (AkT, MAPK) by interacting with cytoplasmic cellular partners, thereby transforming infected cells (Maeda and Fan, 2008; Monot et al., 2015; Maeda et al., 2021). As shown for JSRV, the cellular protein RaLBP1 interacts with JSRV Env to promote AkT/mTOR activation and cell proliferation (Monot et al., 2015). Of particular importance, a YXXM motif located at the C-terminal end of the cytoplasmic tail is involved in cell transformation in numerous cell types and is critical for tumorigenesis *in vivo* (Palmarini et al., 2001; Alberti et al., 2002; Liu et al., 2003; Cousens et al., 2007).

Endogenous retroviruses (ERV), which are closely related to JSRV and ENTVs are present in high copy numbers in the small ruminant genomes. They result from iterative events of integration of retroviral copies into the germ cells, and their vertical transmission to offspring. Endogenous β-retroviruses genetically close to exogenous JSRV and ENTV have been reported in small ruminants (Arnaud et al., 2007; Sistiaga-Poveda and Jugo, 2014; Cumer et al., 2019; Moawad et al., 2024; Verneret et al., 2024) and were initially named enJSRV and enENTV depending on the species from which they were characterized. Their integration into the genome occurred prior to the speciation of *Ovis* and *Capra* genera (Arnaud et al., 2007). The history of their endogenization including the role of JSRV and ENTVs in this process remains to be characterized. In contrast to their exogenous counterparts, the ERV envelope is not oncogenic *in vitro* (Palmarini et al., 2001), due to major differences in the oncogenic region at the end of the TM protein, in particular the absence of the YXXM motif. Since the exogenous viruses at the origin of ERVs are not known, enJSRV and enENTV nomenclature may be misleading. Sheep (sERV) or goat (gERV) ERVs related to oncogenic small ruminant beta retroviruses, focusing on the genome in which they have been described, may be more appropriate and will be used in this article. Recently, we and others have shown that these sERV and gERV are closely related and belong to the same genetic family II-5 (Verneret et al., 2024) or Cap ERV24 (Moawad et al., 2024).

Although known for decades and compared to other animal and human RNA viruses, the genetic diversity of JSRV and ENTVs is only partially known with only a handful of sequences reported, most of which are not associated with clinical status. Only 9 complete sequences of JSRV have been reported originating from Africa, North America, Europe and Asia (York et al., 1992; Palmarini et al., 1999; DeMartini et al., 2001). Fewer than 50 complete genomes of ENTV have been reported, originating from Canada and Europe for ENTV-1 (Cousens et al., 1999; Walsh et al., 2010) or from Europe and Asia for ENTV-2 (He et al., 2017; Ye et al., 2019). Nevertheless, genomic epidemiology is key to understanding the circulation of these oncogenic retroviruses in sheep and goat populations worldwide, and ultimately to genetically tracing pathogenic strains. Specific amplification of the exogenous viruses at the origin of respiratory cancers is even more challenging due to the presence of the sERV and gERV which are highly related to JSRV and ENTVs and should be carefully considered.

In France, lung and nasal cancers induced by oncogenic β-retroviruses are present in small ruminants but are often unrecognized, although some clinical signs such as clear and abundant secretions should alert (Leroux and Turpin, 2021; Devos et al., 2023). These virus-induced cancers are enzootic in France, but the frequency of clinical expression is variable, with flocks with low (sporadic) to high (outbreaks) incidence of cancers (Leroux and Turpin, 2021; Devos et al., 2023). In the latter case, as we have observed on several occasions, the high incidence of cancer can lead to the loss of the flock through uncontrollable mortality directly attributable to cancer or culling. This raises the issue of the reintroduction of retrovirus-free animals, which is difficult to achieve without routine detection and an established control program, and thus represents major threat to livestock production. Our ongoing work clearly demonstrates that ENTVs and JSRV are actively circulating in flocks but have never been genetically characterized.

In the present study, we addressed the diversity of the circulating strains of JSRV and ENTVs in France by developing highly specific PCRs targeting exogenous β-retroviruses in the LTR non-coding region that controls virus expression, in the *env* oncogene and by amplifying and sequencing the near full-length provirus. We have completed the description of fifty-three strains of JSRV and ENTVs and have observed JSRV genetic signatures that may be associated with disease severity.

## 2 Materials and methods

### 2.1 Biological samples

Twenty-four ENTV-induced nasal tumors and twenty-nine JSRV induced lung tumors collected between 2003-2023 originated from 29 different flocks or areas in France have been selected from our biobank (Table 1). Based on information provided by Vets and breeders, the cancer frequency was graded as “low” (sporadic cases over a period of 1–2-years), “high” (multiple cases per year) or “unknown” (no information about the evolution of the number of cases). Cell lines IDO5 (dermal fibroblasts) obtained from Institut Mérieux, Lyon, France and TIGEF (T-immortalized goat embryonic fibroblasts) (Da Silva Teixeira et al., 1997) were used as non-infected controls. Genomic DNAs were extracted from tissues and cells using the “Quick-DNA midiprep plus” kit (Zymo), after a mechanical grinding step (FastPrep device, MP Biomedicals) for tumor tissues.

**Table 1.** Origin of the JSRVs and ENTV. Viruses are referred by their ID #. “+”: sequenced; “0”: not sequenced (no amplification or not enough material available); ”nFL”: near full-length, ID = identification number, “Department” refers to the French departments (identified by their two-digit numbers) the viruses originated from. The cancer frequency was graded as “low” corresponds to sporadic cases over a period of 1–2-years; “High” corresponding to multiple (often several dozen per year) occurring during the same period of time which may lead to the elimination of the flock by premature death of the animals or slaughter. “Unknown” corresponds to flocks for which no information about the evolution of the number of cases was available.

|  | Virus |  |  |  | Flock information |  |  |  |
| --- | --- | --- | --- | --- | --- | --- | --- | --- |
|  | ID # | env | LTR | nFL | ID # | Department | Cancer frequency | Year |
| ENTV-1 | 4403 | 1 | 1 | 1 | FR28 | 27 | High | 2023 |
|  | 4489 | 1 | 0 | 0 | FR29 | 37 | Unknown | 2023 |
|  | 4491 | 1 | 1 | 1 | FR30 | 56 | Unknown | 2023 |
| ENTV-2 | 1039 | 1 | 1 | 1 | FR2 | 23 | Low | 2005 |
|  | 1040 | 1 | 0 | 0 | FR2 | 23 | Low | 2005 |
|  | 1216 | 1 | 0 | 1 | FR4 | 36 | Low | 2007 |
|  | 1255 | 1 | 0 | 0 | FR7 | 79 | Low | 2007 |
|  | 1256 | 1 | 0 | 0 | FR7 | 79 | Low | 2007 |
|  | 1259 | 1 | 0 | 0 | FR7 | 79 | Low | 2007 |
|  | 2853 | 1 | 1 | 0 | FR1 | 74 | High | 2016 |
|  | 2855 | 1 | 1 | 1 | FR1 | 74 | High | 2016 |
|  | 2860 | 1 | 0 | 0 | FR1 | 74 | High | 2016 |
|  | 2876 | 1 | 1 | 1 | FR1 | 74 | High | 2016 |
|  | 2980 | 1 | 0 | 0 | FR1 | 74 | High | 2016 |
|  | 3313 | 1 | 0 | 0 | FR9 | 79 | Low | 2019 |
|  | 3384 | 1 | 1 | 1 | FR8 | NA | Unknown | 2020 |
|  | 3719 | 1 | 0 | 0 | FR10 | 46 | High | 2021 |
|  | 3720 | 1 | 0 | 0 | FR10 | 46 | High | 2021 |
|  | 3817 | 1 | 0 | 0 | FR10 | 46 | High | 2021 |
|  | 3824 | 1 | 1 | 1 | FR6 | 46 | Unknown | 2021 |
|  | 3955 | 1 | 1 | 1 | FR10 | 46 | High | 2021 |
|  | 4196 | 1 | 1 | 1 | FR6 | 46 | Unknown | 2022 |
|  | 4322 | 1 | 1 | 0 | FR5 | 42 | High | 2022 |
|  | 4407 | 1 | 1 | 0 | FR3 | 46 | Unknown | 2023 |
| JSRV | 985 | 1 | 1 | 1 | FR27 | 64 | Low | 2003 |
|  | 987 | 1 | 1 | 1 | FR12 | 64 | Low | 2003 |
|  | 989 | 1 | 1 | 1 | FR21 | 64 | Low | 2003 |
|  | 1000 | 1 | 1 | 1 | FR20 | 64 | Low | 2003 |
|  | 1006 | 1 | 1 | 1 | FR20 | 64 | Low | 2004 |
|  | 1024 | 1 | 0 | 0 | FR14 | 64 | Low | 2005 |
|  | 1025 | 1 | 0 | 0 | FR14 | 64 | Low | 2005 |
|  | 1037 | 1 | 1 | 1 | FR18 | 64 | Low | 2005 |
|  | 1167 | 1 | 1 | 1 | FR13 | 64 | Low | 2006 |
|  | 1296 | 1 | 1 | 1 | FR19 | 64 | Low | 2008 |
|  | 1298 | 1 | 1 | 1 | FR22 | 64 | Low | 2008 |
|  | 1466 | 1 | 1 | 1 | FR24 | 64 | Low | 2008 |
|  | 1468 | 1 | 1 | 1 | FR18 | 64 | Low | 2009 |
|  | 1481 | 1 | 1 | 1 | FR25 | 64 | Unknown | 2009 |
|  | 1506 | 1 | 0 | 0 | FR25 | 64 | Unknown | 2009 |
|  | 1751 | 1 | 1 | 1 | FR26 | 64 | Low | 2011 |
|  | 1762 | 1 | 1 | 1 | FR26 | 64 | Low | 2011 |
|  | 2054 | 1 | 1 | 1 | FR15 | 46 | High | 2012 |
|  | 2055 | 1 | 0 | 0 | FR15 | 46 | High | 2012 |
|  | 2056 | 1 | 0 | 0 | FR15 | 46 | High | 2012 |
|  | 2332 | 1 | 1 | 1 | FR15 | 46 | High | 2012 |
|  | 2333 | 1 | 1 | 1 | FR15 | 46 | High | 2012 |
|  | 2334 | 1 | 1 | 1 | FR15 | 46 | High | 2012 |
|  | 2339 | 1 | 0 | 0 | FR17 | 46 | Low | 2012 |
|  | 2369 | 1 | 1 | 1 | FR15 | 46 | High | 2013 |
|  | 2529 | 1 | 1 | 1 | FR17 | 46 | Low | 2014 |
|  | 2586 | 1 | 1 | 1 | FR16 | 46 | Low | 2014 |
|  | 2780 | 1 | 1 | 1 | FR15 | 46 | High | 2016 |
|  | 3431 | 1 | 0 | 0 | FR23 | 25 | High | 2020 |

### 2.2 Amplification and sanger sequencing of LTR and *env*

The LTR and *env* regions of JSRV, ENTV-1 and ENTV-2 were amplified by PCR (Table 2) using the high-fidelity DNA polymerase “PrimeStar GXL” (Takara), with primers targeting exogenous β-retroviruses. The specificity of the primers was tested first *in silico* by BLASTn analysis on sERV and gERV (see <u>“Sequence used for the phylogenetic analysis”</u> section of the Materials and methods), and on *Ovis aries* (Ramb2.0) and *Capra hircus* (ARS1.2) genomes. The primers used to amplify JSRV and ENTVs were iteratively improved throughout the study based on the sequenced strains (Table 2, <u>Figure S1</u>). The specificity of the produced amplicons was confirmed by the absence of amplification from sheep (IDO5) and goat (TIGEF) DNA (<u>Figure S4</u>).

**Table 2.**
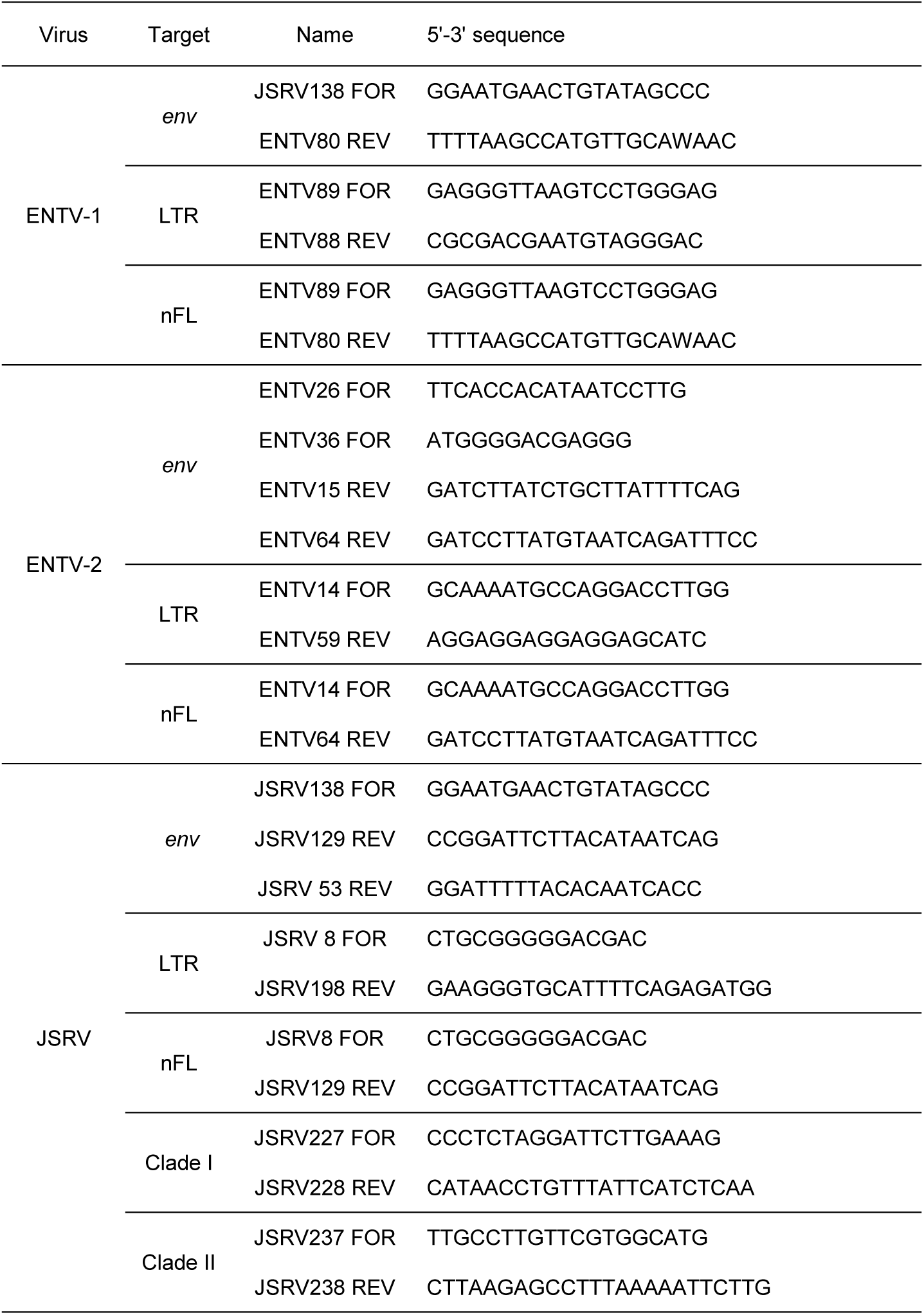
Primers used to amplify LTR, *env* or the near full-length (nFL) proviral genomes.

Sanger sequencing was performed on the *env* and LTR amplicons with a minimum coverage of 2X (Sequencing primers available in <u>Table S1</u>). Contig sequences were generated and mapped to the reference sequences of JSRV (M80216.1 and AF357971.1), ENTV-1 (NC007015.1) and ENTV-2 (NC_004994.2) using the “Geneious Prime 2023.2.1” software. The complete U3-R-U5 LTRs were reconstructed by assembling sequences from the 5’ LTR amplicons and *env* amplicons carrying part of the 3’ LTR.

### 2.3 Whole provirus amplification and genetic analysis

Near full-length provirus amplicons were generated by PCR using “Prime STAR GXL” (Takara) with primers located in the U3 region of the 5’ and 3’ LTRs (<u>Figure S1</u>) and purified using “1X AmpureXP beads” (Beckman Coulter). ONT (Oxford nanopore technologies) libraries were indexed and prepared using the “Native Barcoding kit” (SQK-NBD114.24) and sequenced on MinION Mk1c with R10.4.1 flowcells. All libraries were base-called using the High Accuracy Basecalling (HAC) model of Dorado v0.4.3(Oxford Nanopore PLC, 2023). After demultiplexing, reads were mapped to JSRV (M80216.1 and AF357971.1), ENTV-1 (NC007015.1) and ENTV-2 (NC_004994.2) reference sequences using minimap2 v2.26 (Li, 2018). Consensus sequences were generated from mapped reads of at least 5000 bp using “samtools consensus v1.18” (Danecek et al., 2021) with default settings for HAC basecall nanopore reads. Sequencing data generated in this study were deposited in the NCBI BioSample SRA database (project number PRJNA1103156).

Nucleotide and amino acid alignments were generated with by the "MAFFT Alignment v7.490" tool (Katoh and Standley, 2013). JSRV and ENTVs’ ORFs were determined by alignment with previously published sequences. As other retroviruses JSRV and ENTVs, Pro and Pol are suspected to be produced by a rimosomal frameshift. Thus, we determine the start of Pro and Pol as producing the longest ORF possible (York et al., 1992; Cousens et al., 1999; Walsh et al., 2010).

JSRV variability was determined from the nucleotide and predicted amino acid alignment matrix of each ENTVs and JSRV ORFs. The whole genome identity variation along the JSRV and ENTV-2 genomes were calculated in “Geneious Prime 2023.2.1” on the reconstituted full-length nucleotide sequences alignment then plotted in a window of 150 nt and steps of 10 nt. The JSRV amino acid pairwise identity variation was calculated on individual alignment for each ORF and plotted in a window of 10 aa and steps of 2 aa. Only one sequence per flock was conserved for the JSRV and ENTVs diversity assessment with the exception of flock FR 26 where JSRV clade I and clade II were both detected (1751 and 1762 respectively).

### 2.4 Transcriptomic analysis of JSRV infected lambs

Transcriptomic data generated by Karagianni et al (Karagianni et al., 2019) were re-analyzed to discriminate between mRNA originated from JSRV or from sERV in JSRV-infected lambs. Three sets of lung RNA-seq data were used, two from JSRV-infected lambs (ERR2683948 & ERR2683946) and one from a non-infected lamb (ERR2683952). The search for JSRV or sERV specific transcription signal was performed using BBduk v38.95 (BBMap, 2023), with specific JSRV and sERV k-mers (TCTGATTATGTAAGAATCCGG and GAGAGTTTTAATACATAAAAA respectively) allowing two mismatches. The k-mers were designed from JSRV and sERV available in the public databases. Reads containing these k-mers were then mapped to the reference JSRV strain (AF105220.1) or sERV (DQ838493.1) as a validation step using BWA-mem v0.7.17 (Li and Durbin, 2009) and duplicate reads were removed using the “PICARD Markduplicates tool v4.0.11” (GATK) (Picard Tools - By Broad Institute).

### 2.5 Rapid genotyping of circulating clade I and II JSRVs

For JSRV clade-specific PCRs, primers were designed in the U3 region of the LTR for JSRV clade I and in the *env*-U3 region for clade II (Table 2). The mammalian *thyroglobulin* gene was used as a control to confirm the presence of amplifiable DNA.

### 2.6 Phylogenetic analysis and genetic diversity

LTR, *env* and nFL (near full-length) proviral sequence alignments were performed using "MAFFT Alignment v7.490" (Katoh and Standley, 2013) in “Geneious Prime 2023.2.1”. Variability values were determined by the distance matrix of the alignment file, calculated as the percentage of bases differing between two sequences. Pairwise identity rate for multiple sequences were directly calculated from the alignment. Phylogenetic maximum likelihood trees in the *env* and LTR regions were constructed using the "IQ-TREE v1.5.3" tool (Nguyen et al., 2015) using an ultrafast bootstrap parameter of 10,000 iterations. Identical sequences from a same flock were removed for the *env* and LTR phylogenetic analysis. A minimal branch support of 80 was considered as a bootstrap threshold to define a phylogenetic clade.

### 2.7 Transcription factor binding site prediction

Consensus sequences were generated with “Geneious Prime 2023.2.1” on the French JSRV clade I or clade II LTR sequence alignment with a 75% threshold (Supplemental material JSRV_LTR_consensus.fasta). All the French LTR sequences were used for the consensus generation according to their classification in JSRV clade I or clade II (Table 1). Potential binding sites for transcription factors in JSRV and ENTV LTR were identified with the "EMBOSS 6.5.7 tool tfscan" on the eukaryotic transcription factor "Transfac" database (Matys et al., 2006) using both the “Vertebrate” and “Other” databases. The position of each transcription factor binding site was determined on the LTR clade A and clade B consensus alignment.

### 2.8 Nucleotide sequence accession numbers

The Env coding sequences obtained in the present study have been deposited in GenBank with the following accession numbers: PP575093(#4403), PP719694(#4489), PP575094(#4491), PP575073(#1039), PP575074(#1040), PP575075(#1216), PP575076(#1255), PP575077(#1256), PP575078(#1259), PP575079(#2853), PP575080(#2855), PP575081(#2860), PP575082(#2876), PP575083(#2980), PP575084(#3313), PP575085(#3384), PP575086(#3719), PP719695(#3720), PP575087(#3817), PP575088(#3824), PP575089(#3955), PP575090(#4196), PP575091(#4322), PP575092(#4407), PP575010(#985), PP575009(#987), PP575011(#989), PP575012(#1000), PP575013(#1006), PP575014(#1024), PP575015(#1025), PP575016(#1037), PP575017(#1167), PP575018(#1296), PP575019(#1298), PP575020(#1466), PP575021(#1468), PP575022(#1481), PP575023(#1506), PP575024(#1751), PP575025(#1762), PP575026(#2054), PP575027(#2055), PP575028(#2056), PP575029(#2332), PP575030(#2333), PP575031(#2334), PP575032(#2339), PP575033(#2369), PP575034(#2529), PP575035(#2586), PP575036(#2780), PP575037(#3431). Reconstructed LTR are available on Genbank with the accession numbers: PP575070(#4403), PP575071(#4489), PP575072(#4491), PP575060(#1039), PP575061(#2853), PP575062(#2855), PP575063(#2876), PP575064(#3384), PP575065(#3824), PP575066(#3955), PP575067(#4196), PP575068(#4322), PP575069(#4407), PP575038(#985), PP575039(#987), PP575040(#989), PP575041(#1000), PP575042(#1006), PP575043(#1037), PP575044(#1167), PP575045(#1296), PP575046(#1298), PP575047(#1466), PP575048(#1468), PP575049(#1481), PP575050(#1751), PP575051(#1762), PP575052(#2054), PP575053(#2332), PP575054(#2333), PP575055(#2334), PP575056(#2369), PP575057(#2529), PP575058(#2586), PP575059(#2780). Proviral consensus sequences are available on Genbank with the accession numbers: PP669280(#4403), PP707036(#4491), PP707056(#1039), PP707059(#1216), PP707060(#2855), PP707061(#2876), PP669281(#3824), PP707057(#3955), PP707058(#4196), PP707040(#985), PP707043(#987), PP707047(#989), PP707039(#1000), PP707041(#1037), PP707044(#1167), PP707037(#1296), PP707046(#1298), PP707038(#1466), PP707042(#1468), PP646154(#1481), PP707055(#1751), PP707045(#1762), PP707050(#2054), PP707051(#2332), PP707054(#2333), PP707053(#2334), PP646155(#2369), PP707052(#2529), PP707049(#2586), PP707048(#2780).

### 2.9 Sequence used for the phylogenetic analysis

Publicly available sequences used in the study were: KU258881.1, KU258882.1, KU258883.1, KU258884.1, KU258885.1, KU258886.1, MN564749.1, MN564750.1, MN564751.1 for gERV (Taxonomy ID: 2762664) ; FJ744146.1, FJ744147.1, FJ744148.1, FJ744149.1, FJ744150.1, GU292314.1, GU292315.1, GU292316.1, GU292317.1, GU292318.1, KC189895.1, NC007015.1, for ENTV-1 (TaxID: 69576) and KU179192.1, KU258870.1, KU258870.1, KU258871.1, KU258872.1, KU258873.1, KU258874.1, KU258875.1, KU258876.1, KU258877.1, KU258878.1, KU258879.1, KU258880.1, LC570918.1, LC762616.1, LC762617.1, MF033071.1, MK164396.1, MK164400.1, MK210250.1, MK559457.1, MT254061.1, MT254062.1, MT254063.1, MT254064.1, MT598195.1, NC_004994.2, ON843769.1, OQ989633.1, OR024676.1, OR965522.1, PP130116.1, PP130117.1, PP130118.1 for ENTV-2 (Taxonomy ID: 2913605, 2913605 and 2584748); AF136224.1, AF136225.1, AF153615.1, EF680296.1, EF680297.1, EF680298.1, EF680299.1, EF680300.1, EF680301.1, EF680302.1, EF680303.1, EF680304.1, EF680305.1, EF680306.1, EF680307.1, EF680308.1, EF680309.1, EF680310.1, EF680311.1, EF680312.1, EF680313.1, EF680314.1, EF680315.1, EF680316.1, EF680317.1, EF680318.1, JQ995521.1 for sERV (Taxonomy ID: 9940 (*ovis aries)* defined as “enJSRV”); and A27950.1, AF105220.1, AF357971.1, CQ964469.1, DQ838494.1, KP691837.1, KY041630.1, M80216.1, MN161849.1, MZ931278.1, ON204347.1, OR729406.1, Y18302.1, Y18303.1, Y18304.1, Y18305.1 for JSRV (TaxID: 11746 and 47898).

## 3 Results

### 3.1 Sequencing strategy to specifically amplify exogenous JSRV and ENTVs

While sEVR or gERV are expressed in a variety of tissues in virus-free sheep, we questioned their expression in infected tissues based on transcriptomic dataset (Karagianni et al., 2019) from experimentally JSRV-infected lambs. We demonstrated the presence of both endogenous and exogenous RNAs in the lung (Table 3). This invalidates approaches that use RNA as a safe matrix to specifically detect exogenous ß-retroviruses. Phylogenetic trees were reconstructed from 87 complete sequences tagged as JSRV, ENTV-1, ENTV-2, sERV and gERV found in public databases, (Figure 1, <u>Figure S3</u>). The tree based on complete genomes (Figure 1A) showed that several sequences identified as exogenous JSRV (DQ838494.1 CH, MZ931278 CH), or exogenous ENTVs (KU258870.1 CH, MK164396.1 CH) clearly clustered with ERVs, highlighting the challenge of specifically detecting exogenous oncogenic retroviruses.

**Figure 1:**
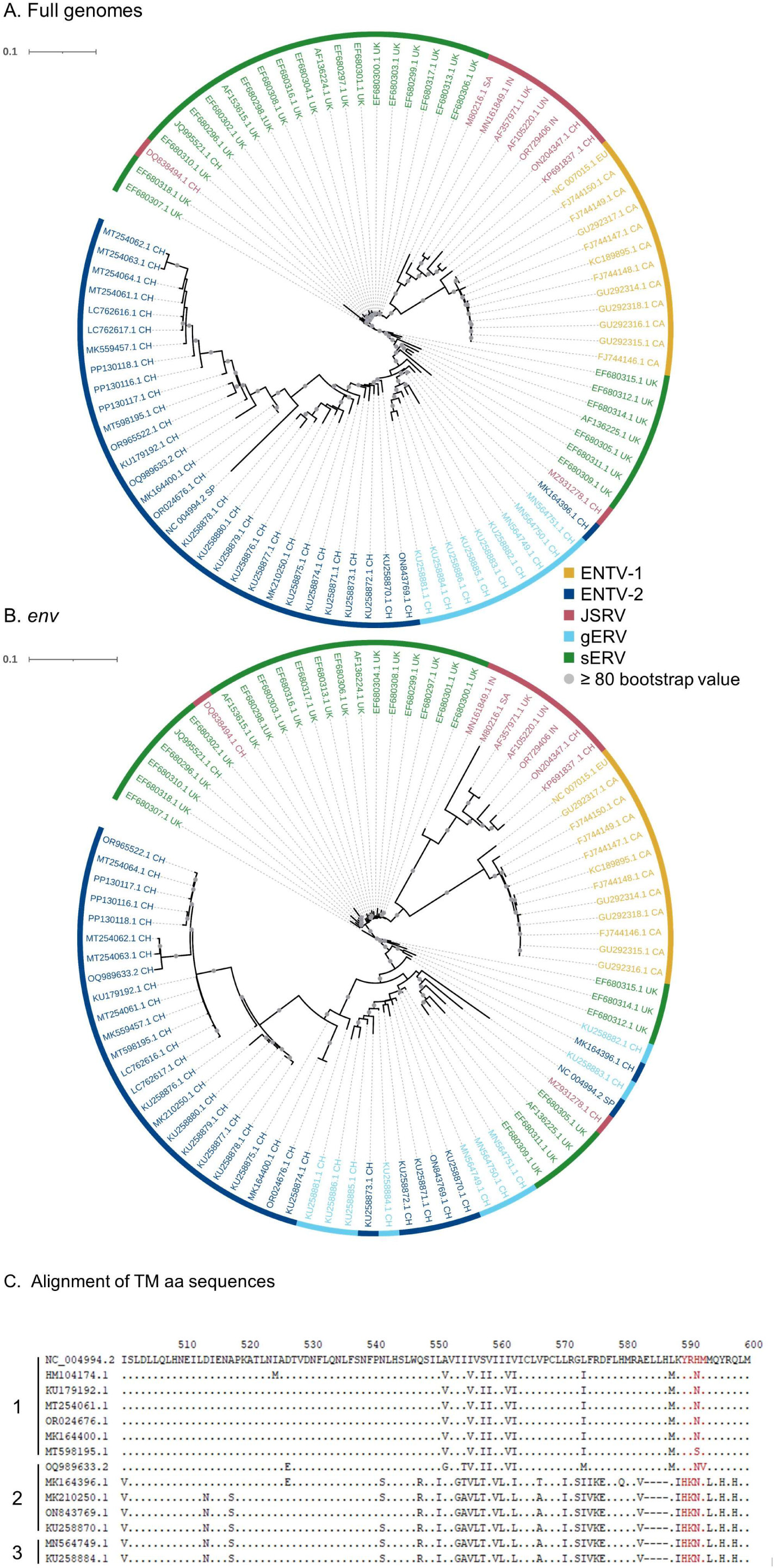
Complexity of distinguishing between the exogenous retroviruses and their endogenous counterparts in small ruminants. Phylogenetic analysis of ENTV-1, ENTV-2, JSRV sequences and their related endogenous sequences from sheep and goats, available in public databases. Maximum likelihood tree with an ultrafast bootstrap parameter of 10,000 iterations using EF680307.1 (enJSRV10) as the root sequence for A) the full-length proviral sequence and B) the *env* region. The last two letters of each sequence define the geographic origin. CH: China, UK: United Kingdom, CA: Canada, IN: India, SA: South Africa, SP: Spain, EU: Europe, UN: Unknown. Bootstrap values greater than 80% are represented by a grey dot and the branch length is indicative of the number of substitutions per site. The scale bar indicates the number of nucleotide substitutions per site. C) Alignment of deduced amino acid sequences of the ENTV-2 and gERV Env TM region. The position of the YXXM motif, the molecular marker for the transforming exogenous strains of JSRV, ENTV-1 and ENTV-2, is indicated in red. “1” defines the sequences with ENTV-2 as a taxonomic assignment in GenBank (Taxonomy ID: 239365, 2913605 and 2584748) and with a complete YXXM motif. “2” defines the sequences with ENTV-2 as a taxonomic assignment in GenBank but without the YXXM motif. ”3” defines the sequences with gERV as a taxonomic assignment in GenBank (Taxonomy ID: 2762664). Dots represent identical amino acids to NC_004994.2 and dashes indicate gaps in the alignment.

**Table 3:** Expression of JSRV and sERV in infected lambs. . Reads including JSRV and sERV discriminating k-mers were counted with Bbduk tool in infected (ERR2683948 and ERR2683946) and non-infected (ERR2683952) lambs (Karagianni et al., 2019) and expressed as total read counts (with or without read duplicates).

|  | Read counts (raw) |  | Read counts (duplicate removal) |  |
| --- | --- | --- | --- | --- |
|  | JSRV | sERV | JSRV | sERV |
| Infected Lamb1 | 1356 | 302 | 241 | 142 |
| Infected Lamb2 | 3158 | 218 | 312 | 218 |
| Non-infected Lamb | 0 | 110 | 0 | 72 |

Since the role of the YXXM motif in cell transformation has been established, we used the presence of the YXXM sequence as a specific signature of exogenous retroviruses. A consensus sERV sequence (Supplementary material: Consensus_enJSRV-BDD.fasta) was generated from published sERV sequences to identify sERV and gERV copies present in 15 *Caprinae* genome assemblies using BLASTn (<u>Figure S2</u>). Of the 667 endogenous sequences with an *env* part encoding the TM region, the majority (626) contained a HKNM and none of them contained the canonical YXXM motif (<u>Figure S2</u>). This confirms the exogenous signature of this short motif. Some sequences identified as exogenous ENTVs in the databases did not contain the YXXM motif (OQ989633.2, MK164396.1, MK210250.1, ON843769.1, KU258870.1), indicating that they were likely amplified and sequenced from an endogenous retrovirus and were excluded from our subsequent JSRV and ENTV primer definition steps (Figure 1C). Specific amplification of exogenous retroviruses was confirmed using genomic DNA from nasal and lung tumors; the absence of amplification from uninfected IDO5 or TIGEF fibroblasts confirmed the exogenous nature of the amplicons with no cross-reaction with related endogenous sequences (<u>Figures S1</u> and <u>S4</u>). Using this rigorous approach, we have developed molecular tools for the detection and amplification of strictly exogenous JSRVs and ENTVs based on the sERV and gERV genomes present in the genomic databases and on a specific signature that confirms their exogenous status.

### 3.2 ENTV-1 and ENTV-2 cluster in French phylogenetic groups

ENTV-1 and ENTV-2 were sequenced from twenty-four (3 from sheep, and 21 from goats respectively) nasal tumors collected from 13 flocks originating from different departments of France (Table 1). We demonstrated the circulation of a major genetic clade of ENTV-2, with sequences having a pairwise identity > 99.5% for both *env* and LTR regions (Figure 2 A and B), except for 3384FR which is the only French sequence found in an outgroup with between 98 and 99% pairwise identity in LTR and *env* with other French ENTV-2 sequences. All *env* sequences generated in this study had the YXXM motif confirming their exogenous nature. French ENTV-2 strains differed in the *env* region from strains reported in Spain (NC_004994.2, 5.5 % nt divergence) or in China (KU258879.1, 16% nt divergence) (Figure 2<u>.A</u>). Based on the LTR analysis, the French and Spanish ENTV-2 strains clustered together (Figure 2<u>.B</u>).

**Figure 2:**
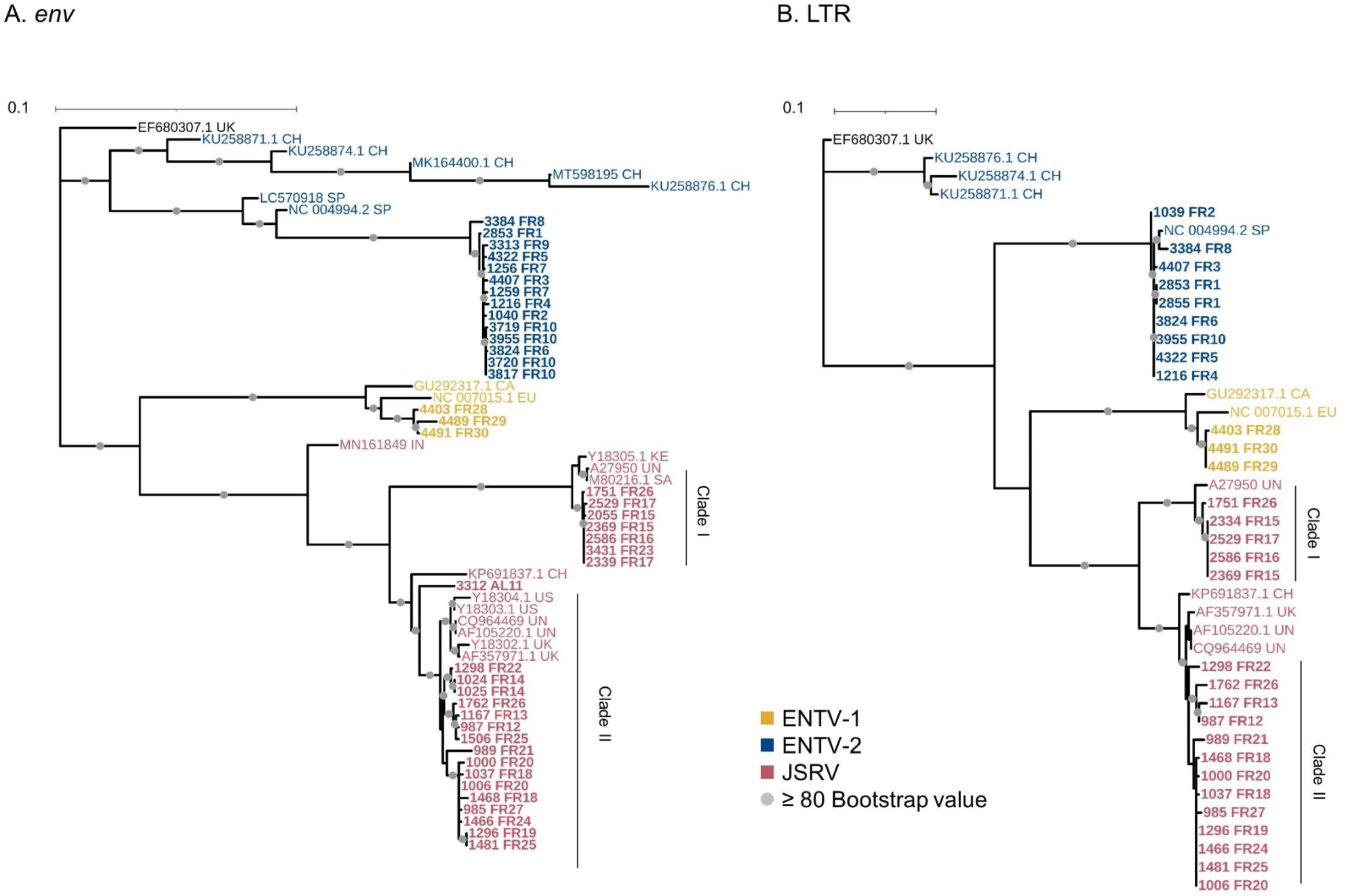
French JSRV and ENTV strains are distributed in distinct phylogenetic clades. A-B) Maximum likelihood tree with an ultrafast bootstrap parameter of 10,000 iterations for A) *env* (1851 bp) and B) reconstituted LTR regions (458 bp). The French sequences described in this study are in bold. CH: China, UK: United Kingdom, CA: Canada, IN: India, SA: South Africa, SP: Spain, EU: Europe, KE: Kenya, US: United States, UN: Unknown. Bootstrap values greater than 80% are represented by a grey dot. The scale bar indicates the number of nucleotide substitutions per site.

The three French ENTV-1 strains clustered together and were distinct from previously reported ENTV-1 in North America (GU292317.1) and Europe (NC007015.1). It should be noted that limited ENTV-1 sequences are available in the databases. Pairwise identity between French ENTV-1 strains was >98.9 % for both LTR and *env* based on the alignment matrix used for the phylogenetic analysis (not shown).

### 3.3 Two distinct JSRV clades are circulating in France

Phylogenetic reconstruction based on *env* and LTR sequences obtained from JSRV present in 29 lung cancers revealed the circulation of at least two distinct clades I and II in the *env* and LTR phylogenetic analysis (Figure 2<u>.A-B</u>). Clade I was genetically related to the South African JSRV reference (M80216.1), whereas clade II was more closely related to the UK JSRV reference (AF357971.1). A unique clade per flock was detected in all but one flock. Co-circulation of both clades was detected in flock FR26 for strains 1751 and 1762, sampled on the same day, and classified as clade I and II JSRVs respectively (Table 1 & Figure 2). Within a clade, the nucleotide divergence was very low, with a mean pairwise identity of 98.9% (+/- 0.5%) for clade I, and 99.9% (+/- 0.06%) for clade II in the *env* region. For two sequences found in the same flock (2334 FR15 & 2333 FR15), we observed a large deletion of 10 nucleotides in the U3 region of the LTR just before the TATA box.

### 3.4 JSRV and ENTV strains are stable between and within flocks

To explore the overall diversity across the entire viral genome, a near complete genome (nFL) sequencing strategy based on LTR and *env* targeted sequencing was developed. Amplicons of approximately 7.6 kb, from 2 ENTV-1, 7 ENTV-2, and 26 JSRV strains were obtained and sequenced using Oxford Nanopore technology (Figure <u>S1</u>-<u>S4</u>, Table 1). The nFL proviral sequencing confirmed that the same major strains of JSRV and ENTV were circulating in the sheep and goat flocks. The ENTV strains have a low variability with 0.29% (+/- 0.08) for ENTV-2 and 0.46% for ENTV-1 for the nFL genome with less than 1% variability in the different genes (Figure 3<u>.A</u>, Table 4). JSRV nFL provirus sequencing confirmed the circulation of JSRV clades I and II in France (Figure 3<u>.B</u>), with 3.5 % variability between the 16 fully sequenced strains (Figure 3<u>.B</u>, Table 4). At the amino acid level, variability was observed in different regions of Gag (Figure 4<u>.A</u>); in the phosphoprotein protein 24, p12 and nucleocapsid (NC) proteins. The N-terminal region encoding for the viral protease (PR) is also variable with a predicted potential smaller size of the Pro CDS due to the presence of two premature stop codon for clade II in this region (Figure 4<u>.B</u>). Variability is observed in the integrase protein (INT) with a premature stop codon at the C-terminus for 5 clade I strains, removing 4 aa (Figure 4<u>.C</u>). A polymorphism hotspot was detected in the *env* region encoding for the cytoplasmic tail. In this 68 amino-acid long region, the analysis showed 92.7 % identity compared to over 98 % identity when considering the full Env CDS (Figure 4<u>.D</u>).

**Figure 3:**
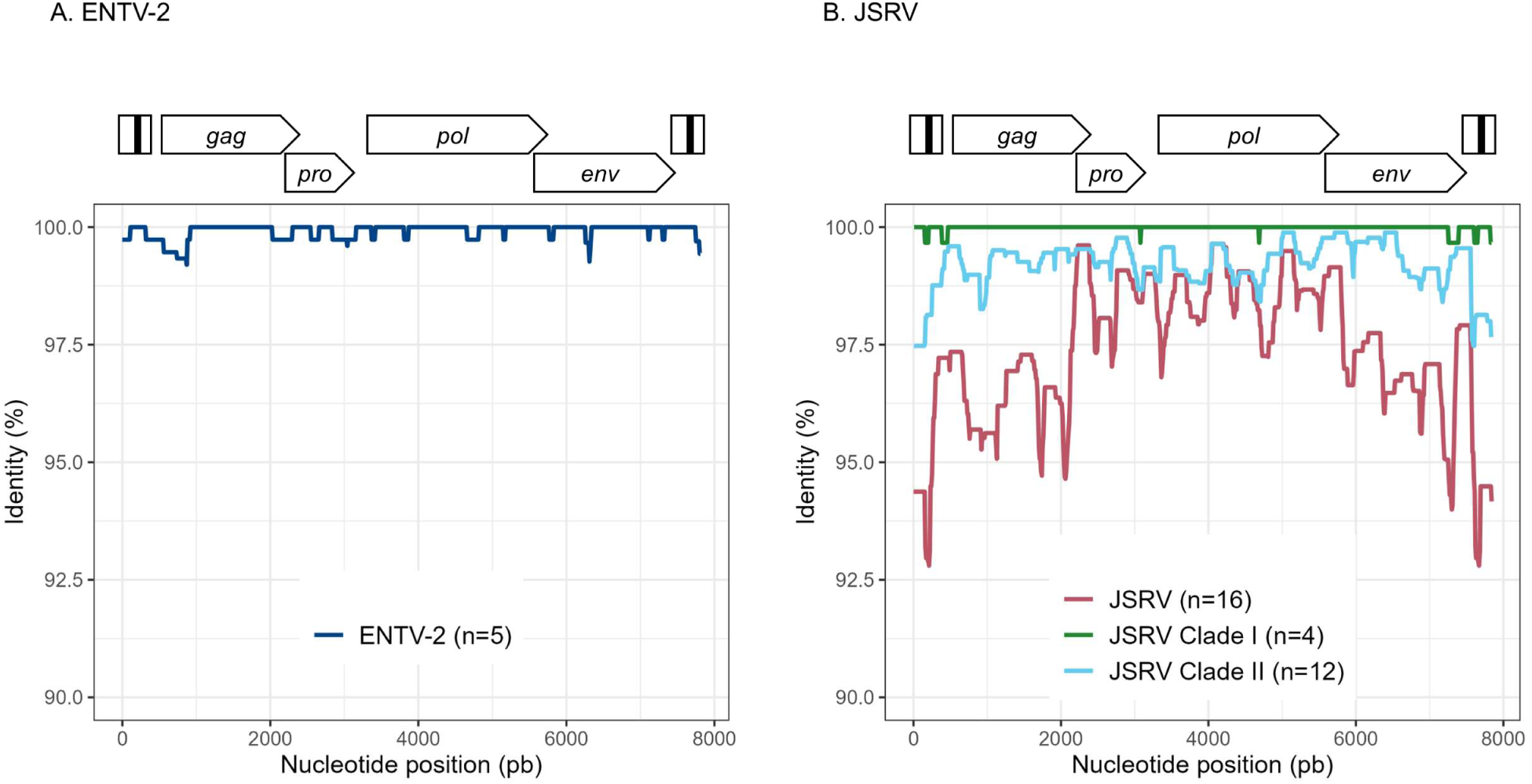
Variability ENTV-2 and JSRV. Nucleotide identity was determined along the reconstituted full-length proviruses of A) ENTV-2 and B) JSRV. Each sequence is from a different flock.

**Figure 4:**
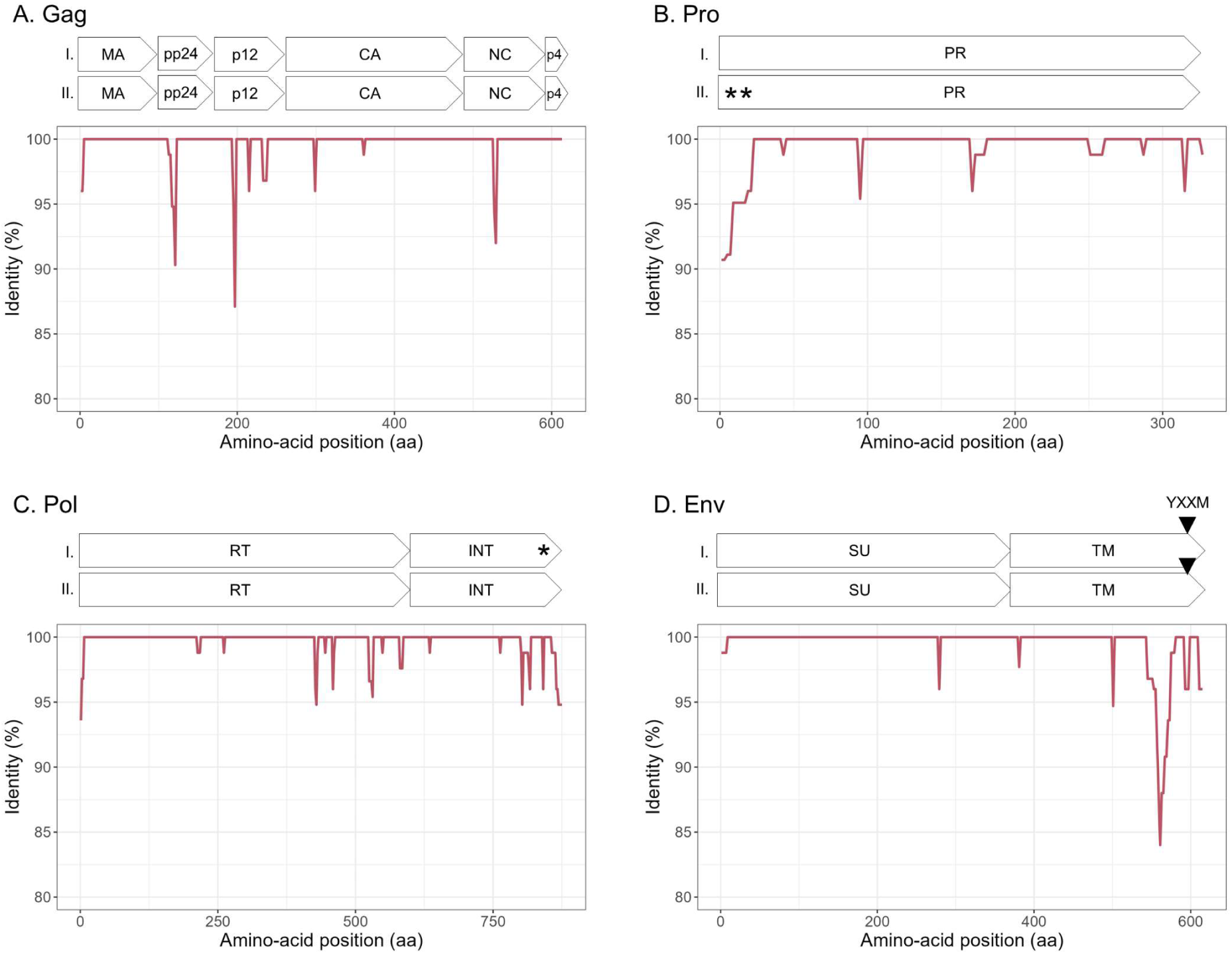
Amino acid sequence diversity of JSRV ORFs. A-D) Calculated pairwise identity in deducted aa along the different ORFs of JSRV calculated with a sliding window of 10 aa and steps of 2 aa. The different protein/domains encoded by the ORF are shown above each graph. The YXXM motif is represented with a grey triangle. Stop codons are indicated by asterisks.

**Table 4:** JSRV and ENTV genetic diversity.

| | | % (± sd) divergence | | | | | | |
| --- | --- | --- | --- | --- | --- | --- | --- | --- |
| Virus | n flocks |  | LTR | *gag* | *pro* | *pol* | *env* | NFL |
| ENTV-1 | 2 | nt | 0.27 | 0.33 | 0.97 | 0.5 | 0.38 | 0.46 |
|  |  | aa | NA | 0.16 | 0.97 | 0.57 | 0.16 | NA |
| ENTV-2 | 5 | nt | 0.32 (± 0.16) | 0.31 (± 0.19) | 0.32 (± 0.20) | 0.21 (± 0.07) | 0.33 (± 0.08) | 0.29 (± 0.08) |
| | | aa | NA | 0.39 (± 0.30) | 0.51 (± 0.47) | 0.22 (± 0.16) | 0.13 (± 0.10) | NA |
| JSRV | 16 | nt | 5.40 (± 4.65) | 4.25 (± 3.94) | 2.53 (± 1.91) | 2.22 (± 1.50) | 4.08 (± 3.80) | 3.56 (± 3.02) |
| | | aa | NA | 1.09 (± 0.92) | 1.02 (± 0.79) | 1.06 (± 0.53) | 1.22 (± 1.14) | NA |

As for three flocks, we collected lung and nasal tumors over several months, we questioned the genetic stability of the viral strains at the origin of cancer over time. Over 1.5 months (flock FR1) or 11 months (flock FR6), ENTV-2 strains were highly stable with 0.03 % and 0.1% nt variability, respectively, for the reconstituted full-length sequence with no variability in the *env* gene. We have collected JSRV tumor samples over 45 months (flock FR15), characterized by a high incidence of lung cancer associated with clade I viruses. Analysis confirmed the high stability of the JSRV genome, with only 0.2 % nt variability across the 6 full-length proviral genomes. Since the time of infection is not known, the mutation rate cannot be estimated. These results highlight the persistence of the same viral strains over time for both JSRV and ENTV-2.

### 3.5 Clade I JSRV is associated with high prevalence of cancers

JSRV-induced lung cancers are generally reported as isolated cases, but flocks with a high incidence of cancer have been regularly observed in France, with a major impact on breeders through increased mortality rate of young adults. Among the 16 flocks from which we collected tumors, three (FR15, FR16, FR23) showed a high incidence of cancer (Table 1) and were associated with the circulation of clade I JSRV (Figure 2A-B). The Env and LTR regions are the most divergent between the clade I and clade II and are involved in the transformation process and proviral expression respectively. The Env sequences of clades I and II showed differences, particularly in the region encoding for the intracytoplasmic tail of TM (Figure 5). The YXXM motif located in the CT region, known to be the major determinant of cell transformation, was present in both clades, but while the clade I virus associated with cancer outbreaks carried a YRTM motif, clade II viruses carried a YRNM motif.

**Figure 5:**
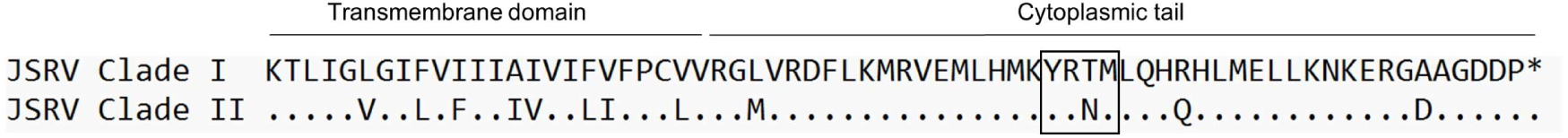
JSRVs clade I and clade II differ in the C-Terminal region of Env. Alignment of the deduced amino acid sequence of 2 representative JSRV clade I (2369 FR) and JSRV clade II (1481 FR) sequences. The localization of the YXXM motif is represented by a black square. The dots represent identical amino acids between the 2 sequences and the asterisk represents the stop codon. YXXM motif starts at the 590^th^ amino acid of JSRV Env.

In the LTR, the differences between JSRV clades I and II were mainly located in the U3 region with 31 mismatches. Prediction of transcription factor binding sites in the LTR using the EMBOSS tool tfscan clearly showed different profiles between the LTRs of the two clades (Figure 6, <u>Table S2</u>). The consensus LTR sequences of JSRV clade I and clade II were aligned, and the position of each transcription factor was indicated from the beginning of U3. A total of 35 transcription factor binding sites were predicted to be common to JSRV clade I and clade II LTRs. Additionally, respectively 11 and 17 transcription factors binding sites were predicted solely in JSRV clade I and clade II LTR. Among them E2F1 (Transcription factor E2F1) at position 105 and CREB (cAMP response element-binding protein) at position 156 were only predicted in JSRV clade I. Specific for JSRV clade II, two TMF (TATA element modulatory factor) at positions 104 and 174 were predicted. The presence of binding sites for the HNF-3, a transcription factor described as modulator of JSRV expression in the lung, at position 127 was confirmed for both clade I and clade II JSRV.

**Figure 6:**
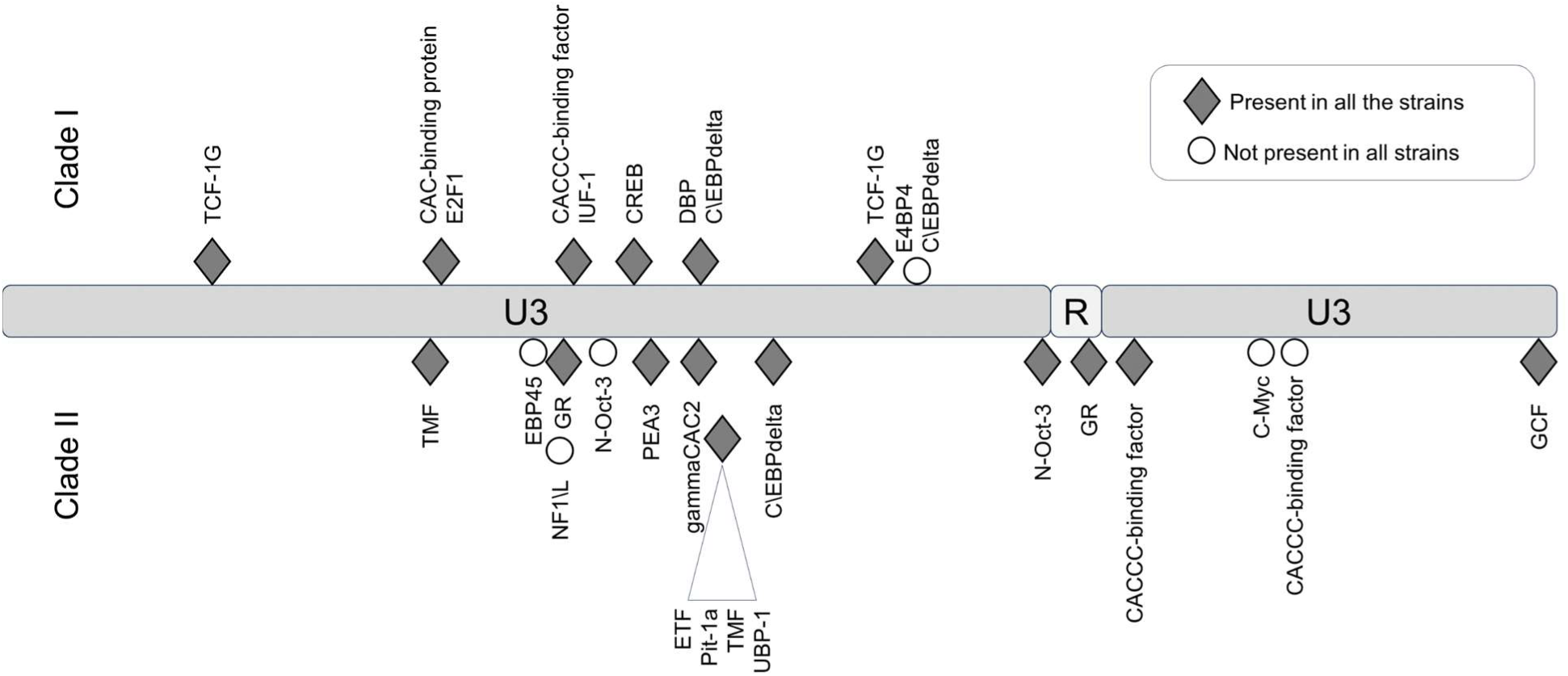
Differential transcription factor binding site prediction on JSRV LTR clade I and clade II. Predicted with the EMBOSS tool TF scan on LTR clade I and clade II sequences, transcription factor binding sites unique to each LTR were presented. Transcription factor binding site prediction was performed on consensus LTR sequences for clade I or clade II. The frequency of each site was tested for both clade using the EMBOSS tool TF scan on each individual LTR sequence (9 sequences for clade I and 13 sequences for clade II). U3: unique 3’; R: Repeat; U5: Unique 5’

### 3.6 Rapid genotyping of JSRV clades

In the absence of routine diagnostics to identify infected sheep, we have developed a PCR-based genotyping tool using discriminating regions between clades I and II (Figure 7). This tool was tested on 6 JSRV strains of known genotype and the amplification of a fragment of 153 bp for clade I and 245 bp for clade II indicated the appropriate genotype.

**Figure 7:**
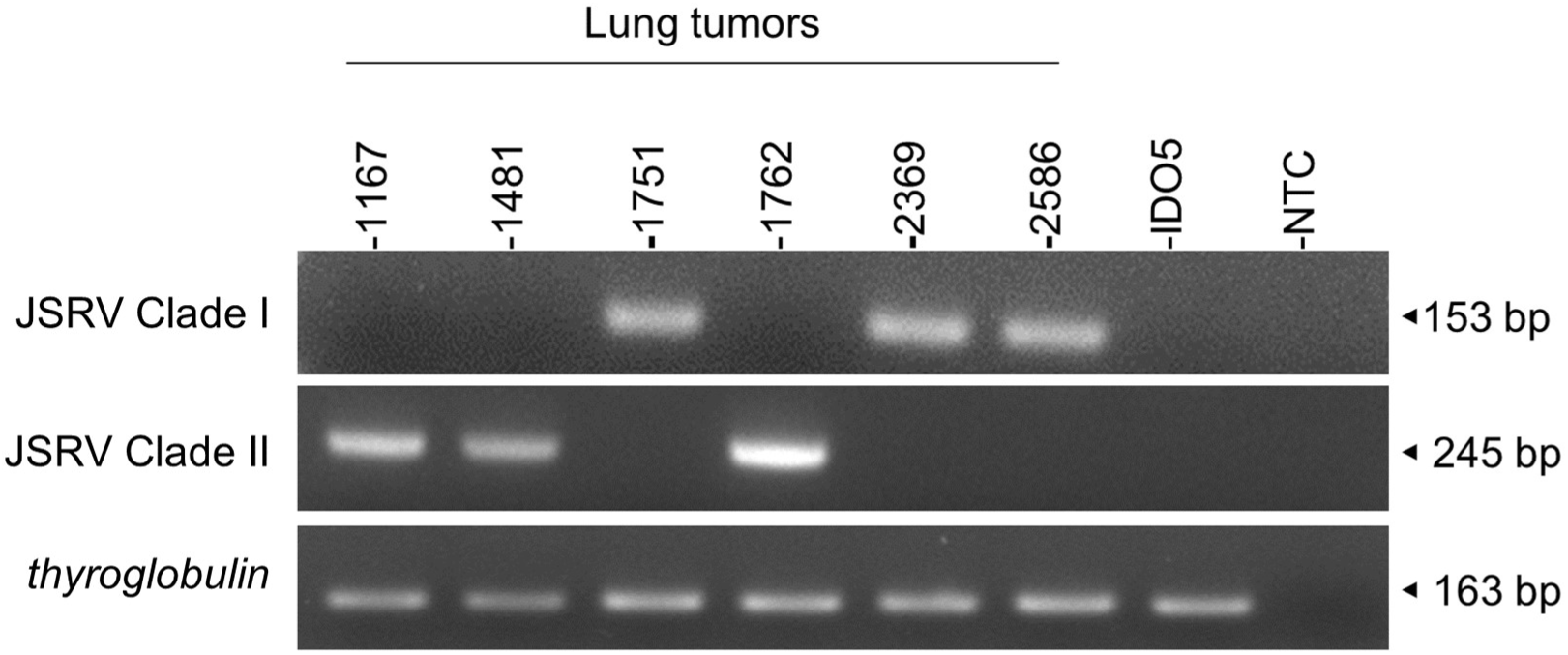
JSRV genotyping by PCR. Agarose gel migration of PCR amplicons corresponding to a region of the JSRV LTR measured at 153 bp for clade I and at 245 bp for clade II. DNA extracted from JSRV-induced tumors of known clade: clade I (1751, 2369 and 2586) and clade II (1169, 1481 and 1762) were used for this proof of concept. IDO5 is an ovine dermal fibroblast cell line used as a JSRV primer specificity control. Thyroglobulin: amplicon corresponding to a region of the *thyroglobulin* gene in the sheep genome, measured at 163 bp.

## 4 Discussion

To determine the diversity of JSRV and ENTV strains circulating in France, we developed both a targeted and near full-length sequencing approach. The detection of RNA from either endogenous or exogenous JSRV in the lung demonstrated the difficulty of performing short-read sequencing on total RNA to reconstruct the full-length genome of exogenous strains. With nearly as much sERV RNA as JSRV RNA, there is a high probability of artificially creating chimeric sequences of both endogenous and exogenous reads in non-discriminating regions. Our targeted sequencing and long read sequencing using primers located in discriminating regions allowed us to be confident in the nature of the strains being sequenced. Furthermore, the detection of endogenous RNA in the infected lung shed light on a possible interaction between endogenous and exogenous RNA/proteins in infected tissues that could modulate the pathogenicity of JSRV as it has already been described *in vitro* with the interaction between endogenous and exogenous Gag (Murcia et al., 2007) or on the possible recombination between endogenous and exogenous strains, as has been observed for feline leukemia virus to form a new pathogenic strain (Erbeck et al., 2021).

Our study has shown that at the national level, a limited number of major strains of JSRV and ENTVs circulate between flocks located in different regions of France, suggesting the introduction of strains that have established and persisted in the animal populations. The genetic stability observed is clearly different from that observed for lentiviruses in livestock, which replicate and mutate rapidly and eventually escape the immune system (Narayan et al., 1977; Leroux et al., 1997; Howe et al., 2002). The minimal diversity observed for ENTVs and JSRV is closer to that reported for HTLV-1 (Human T-cell lymphotropic virus type 1) in humans, with 2-8 % of nt diversity between geographic subtypes, and low (0.5 %) intra-strain variability (Desrames et al., 2014). Compared to other viruses, the number of sequences for JSRV and ENTVs in the public databases is still limited, hindering our understanding of their genetic epidemiology and global circulation. Our efforts to specifically characterize exogenous viruses using tools that exclude sERVs or gERVs, although we have not characterized all the viruses circulating in France, have helped us to define new clades or lineages (Figure 8). French ENTV-2 sequences structured a new lineage, together with the Spanish strain NC_004992.2 (Ortín et al., 2003) which was considered as an outgroup sequence before this work. French ENTV-1 clustered with another European ENTV-1 strain NC007015.1 (Cousens et al., 1999) in a new phylogenetic clade. Interestingly, we found that the French clade I viruses clustered with the JSRV strain first described in South Africa and to our knowledge only reported in Africa (Bai et al., 1999). This may illustrate the circulation of viruses between countries, probably through animal trade.

**Figure 8:**
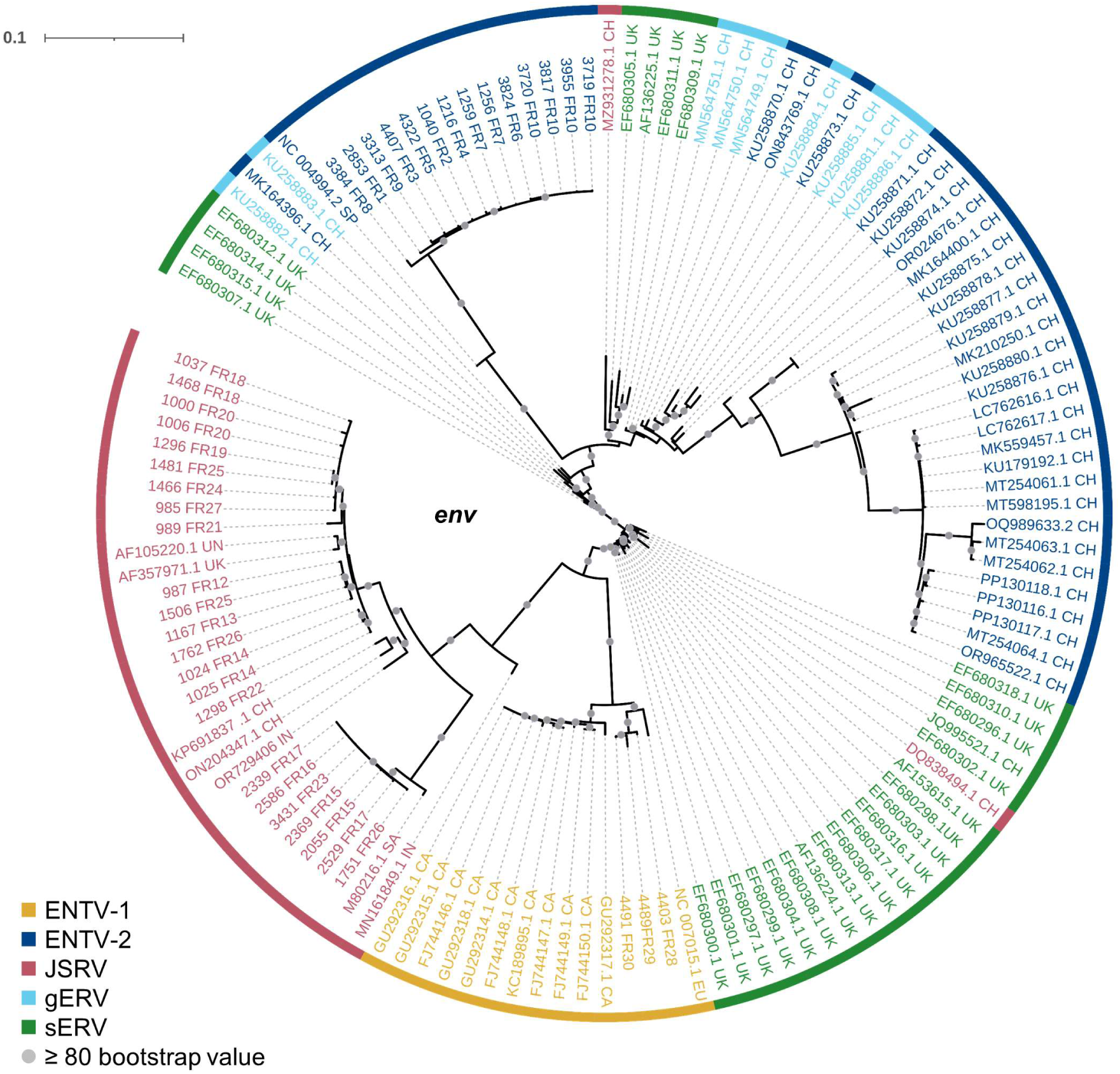
Phylogenetic relationships of the French JSRV and ENTV strains with other endogenous and exogenous small ruminant β-retrovirus. Maximum likelihood tree of the *env* region using 10,000 ultrafast bootstrap repeats with EF680307.1 (enJSRV10) as the rooted sequence. CH: China, UK: United Kingdom, CA: Canada, IN: India, SA: South Africa, SP: Spain, EU: Europe, UN: Unknown. Bootstrap values greater than 80 % are represented by a grey dot and the branch lengths indicate the number of substitutions per site.

We have shown that at the country level and based on our virus collection, the genetic diversity is higher for JSRV than for ENTVs with two distinct clades of JSRV circulating in France. As clade I was mostly associated with flocks with high incidence of cancers, we investigated whether a genetic signature could be detected between clades I and II. Polymorphism analysis along the different JSRV sequences showed a higher degree of polymorphism in Gag and Env. Most differences were observed in the p12 protein and in the nucleocapsid for Gag and in the TM domain for Env. In the p12 protein a L domain involved in JSRV late restriction *in vitro* (Murcia et al., 2007) is found near the polymorphic region. Although no polymorphism is observed on this site, amino acid changes nearby could impact the restriction phenomenon and improve virus spreading in infected individuals. To our knowledge the retroviral nucleocapsid has never been shown to improve pathogenicity in the retrovirus family, but a recent study on coronavirus mutants indicated that point mutation in this protein could improve the infectivity, fitness and virulence of SARS-CoV-2 (Wu et al., 2021). Polymorphism in the transmembrane protein of JSRV is also of interest as it is the major determinant in the transformation process (Palmarini et al., 2001). In the strains analyzed, most of the differences between JSRV clade I and clade II were in the transmembrane part of TM just before the CT domain. This region has been less investigated and its impact on cell transformation remains to be determined. In the YXXM motif, a major determinant of cell transformation, amino acid 592 is found with a threonine/alanine or an asparagine for clades I and II respectively. A glutamine is found at position 598 for clade I while an arginine is present in clade II JSRV. Interestingly, these two amino acids at positions 592 and 598 of the CT domain have been reported to increase cell transformation *in vitro* (Hull and Fan, 2006). In addition for the amino acid at position 592, a change from an asparagine (clade II) to a threonine (clade I) increases the cell transformation in NIH 3T3 cells (Palmarini et al., 2001). We observed differences in the predicted length of the integrase, four amino acids longer for clade I than clade II JSRVs. The C-terminal region of integrase binds to HIV vRNA *via* basic residues during viral particle formation (Elliott et al., 2020), the deletion found in JSRV clade II could interfere with this process and reduce the amount of particles produced. The LTRs of retroviruses are major determinants of viral expression, controlling cell specificity through the pattern of transcription factor binding to the non-coding sequence. Most nucleotide variations are in the viral promoter U3 region with transcription factor binding sites restricted to either the I or II LTR. In U5, the limited differentially predicted restriction factor binding sites could also be detrimental for JSRV expression as a previous study (Palmarini et al., 2000) showed that deletion of this region result in 50 to 80% decrease reporter expression. Altogether modification of LTR activity affect the viral cell cycle like oncogenic envelope protein expression or infectious particle production.

In conclusion, using highly specific sequencing method, we demonstrated that in France with circulation of major strains for ENTV-1 and ENTV-2 as opposed to two clades for JSRV. Genetic signature in the JSRV clade I and clade II could be associated with difference of pathogenicity manifested in flocks by differential cancer incidence. The JSRV polymorphism is mainly located in the TM of Env and the LTR regions involved in cell transformation and viral expression. We also developed a molecular tool to easily follow the circulation of JSRV clades in flocks assess its potential link with disease prevalence. This study will facilitate the monitoring of JSRV and ENTV strains of concern in French flocks. Targeted genetic diagnostic through genomic epidemiology studies could help to reduce the circulation of more pathogenic strains and better understand the dynamics of infection between flocks.

## 6 Author Contributions

BRV, RD, DC, BC realized the experiment, processed the samples and data. BRV and MV performed the analyses. BRV, CL, and JT designed the analysis. BRV and CL wrote the manuscript. CL organized the sample collection with Vets and collected the clinical data. CL and JT conceived the project and secured the fundings. CL, JT and JLC supervised the work.

All participants contributed to the design and implementation of the research, participated to the interpretation of the data and revised the manuscript.

## 7 Acknowledgement

We thank our collaborators from the sheep and goat industry for the collection of biological samples and sharing the clinical information. This work was carried out using the computing facilities of the CC LBBE/PRABI. This study has benefited from the expertise and facilities of the DTAMB and PRABI-AMSB platforms of FR BioEEnViS.

---

**Table S1.**
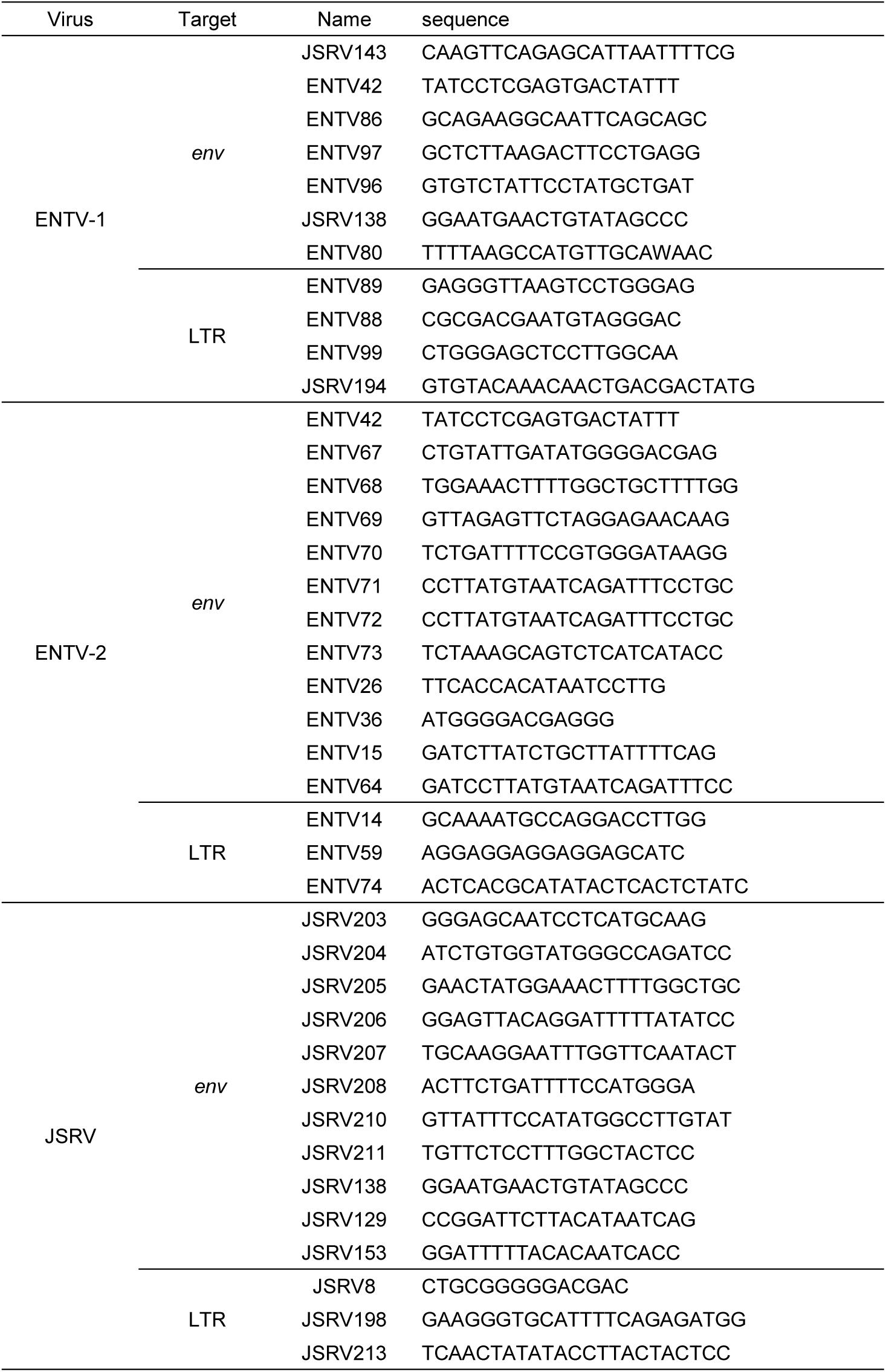
Primers used for the sequencing of JSRV, ENTV-1 and ENTV-2 *env* and LTR amplicons.

**Table S2:** Prediction of transcription factor binding sites in LTR of clades I and II JSRV. sequences were scanned with the tfscan tool to identify transcription factors binding sites on clades I and II LTRs. (+): present, (-): absent. Positions start and end of the transcription factor binding sites are indicated, based on the alignment of the clade I and clade II LTR consensus sequences. The transcription factor frequency is calculated by scanning with the tfscan tool the 22 different LTR sequences of the study (9 LTR-I and 13 LTR-II sequences).

| Name | Transfac<br>Accession | JSRV clade |  | Position<br>Start | Position End | Transcription factor<br>frequency |
| --- | --- | --- | --- | --- | --- | --- |
|  |  | I | II |  |  |  |
| E4BP4 | T01428 | + | + | 2 | 7 | 22/22 |
| ETF | T00270 | + | + | 31 | 35 | 22/22 |
| NF-1/L | T00599 | + | + | 42 | 47 | 22/22 |
| NF-1 | T00537 | + | + | 50 | 55 | 22/22 |
| TCF-1G | T01982 | + | + | 53 | 57 | 22/22 |
| NF-E | T01214 | + | + | 81 | 85 | 22/22 |
| E2F-1 | T01542 | + | + | 82 | 89 | 22/22 |
| GR | T00333 | + | + | 82 | 87 | 22/22 |
| CAC-binding protein | T00076 | + | + | 89 | 93 | 22/22 |
| CACCC-binding factor | T00077 | + | + | 90 | 94 | 22/22 |
| gammaCAC2 | T00075 | + | + | 90 | 94 | 22/22 |
| CTCF | T00172 | + | + | 92 | 96 | 22/22 |
| E1A-F | T01143 | + | + | 123 | 129 | 22/22 |
| HNF-3 | T00371 | + | + | 127 | 133 | 22/22 |
| NF-1/L | T00599 | + | + | 137 | 141 | 22/22 |
| CREB | T00163 | + | + | 161 | 166 | 22/22 |
| Pit-1a | T00691 | + | + | 167 | 173 | 22/22 |
| ETF | T00270 | + | + | 170 | 174 | 22/22 |
| CAC-binding protein | T00076 | + | + | 204 | 208 | 20/22 |
| Pit-1a | T00691 | + | + | 221 | 227 | 22/22 |
| Pit-1a | T00691 | + | + | 223 | 229 | 22/22 |
| ETF | T00270 | + | + | 249 | 253 | 22/22 |
| Pit-1a | T00691 | + | + | 249 | 253 | 22/22 |
| TFIID | T00820 | + | + | 249 | 254 | 22/22 |
| TMF | T00835 | + | + | 249 | 255 | 22/22 |
| NF-1 | T00537 | + | + | 249 | 256 | 22/22 |
| c-Myc | T00140 | + | + | 249 | 256 | 22/22 |
| TMF | T00835 | + | + | 249 | 253 | 22/22 |
| UBP-1 | T00862 | + | + | 249 | 253 | 22/22 |
| ETF | T00270 | + | + | 250 | 256 | 22/22 |
| Pit-1a | T00691 | + | + | 251 | 255 | 22/22 |
| YY1 | T00915 | + | + | 309 | 319 | 22/22 |
| E4BP4 | T01428 | + | + | 338 | 343 | 22/22 |
| E47 | T00207 | + | + | 364 | 370 | 22/22 |
| CAC-binding protein | T00076 | + | + | 391 | 395 | 22/22 |
| E2F-1 | T01542 | + | + | 391 | 395 | 22/22 |
| TCF-1G | T01982 | + | NA | 47 | 51 | 09/09 |
| CAC-binding protein | T00076 | + | NA | 105 | 109 | 09/09 |
| E2F-1 | T01542 | + | NA | 105 | 109 | 09/09 |
| CACCC-binding factor | T00077 | + | NA | 140 | 144 | 09/09 |
| IUF-1 | T00434 | + | NA | 140 | 145 | 09/09 |
| CREB | T00163 | + | NA | 156 | 160 | 09/09 |
| DBP | T00183 | + | NA | 170 | 179 | 09/09 |
| C/EBPdelta | T00583 | + | NA | 171 | 179 | 09/09 |
| TCF-1G | T01982 | + | NA | 228 | 232 | 09/09 |
| E4BP4 | T01428 | + | NA | 233 | 242 | 07/09 |
| C/EBPdelta | T00583 | + | NA | 234 | 242 | 07/09 |
| TMF | T00835 | NA | + | 104 | 109 | 13/13 |
| EBP45 | T01107 | NA | + | 128 | 136 | 12/13 |
| NF-1/L | T00599 | NA | + | 136 | 141 | 12/13 |
| GR | T00333 | NA | + | 137 | 142 | 13/13 |
| N-Oct-3 | T01873 | NA | + | 148 | 156 | 10/13 |
| PEA3 | T00686 | NA | + | 160 | 165 | 13/13 |
| gammaCAC2 | T00075 | NA | + | 172 | 176 | 13/13 |
| ETF | T00270 | NA | + | 174 | 178 | 13/13 |
| Pit-1a | T00691 | NA | + | 174 | 178 | 13/13 |
| TMF | T00835 | NA | + | 174 | 178 | 13/13 |
| UBP-1 | T00862 | NA | + | 174 | 178 | 13/13 |
| C/EBPdelta | T00583 | NA | + | 190 | 198 | 13/13 |
| N-Oct-3 | T01873 | NA | + | 260 | 268 | 13/13 |
| GR | T00333 | NA | + | 273 | 277 | 13/13 |
| CACCC-binding factor | T00077 | NA | + | 285 | 289 | 13/13 |
| c-Myc | T00143 | NA | + | 323 | 329 | 11/13 |
| CACCC-binding factor | T00077 | NA | + | 328 | 332 | 11/13 |
| GCF | T00320 | NA | + | 390 | 396 | 13/13 |

**Figure S1:**
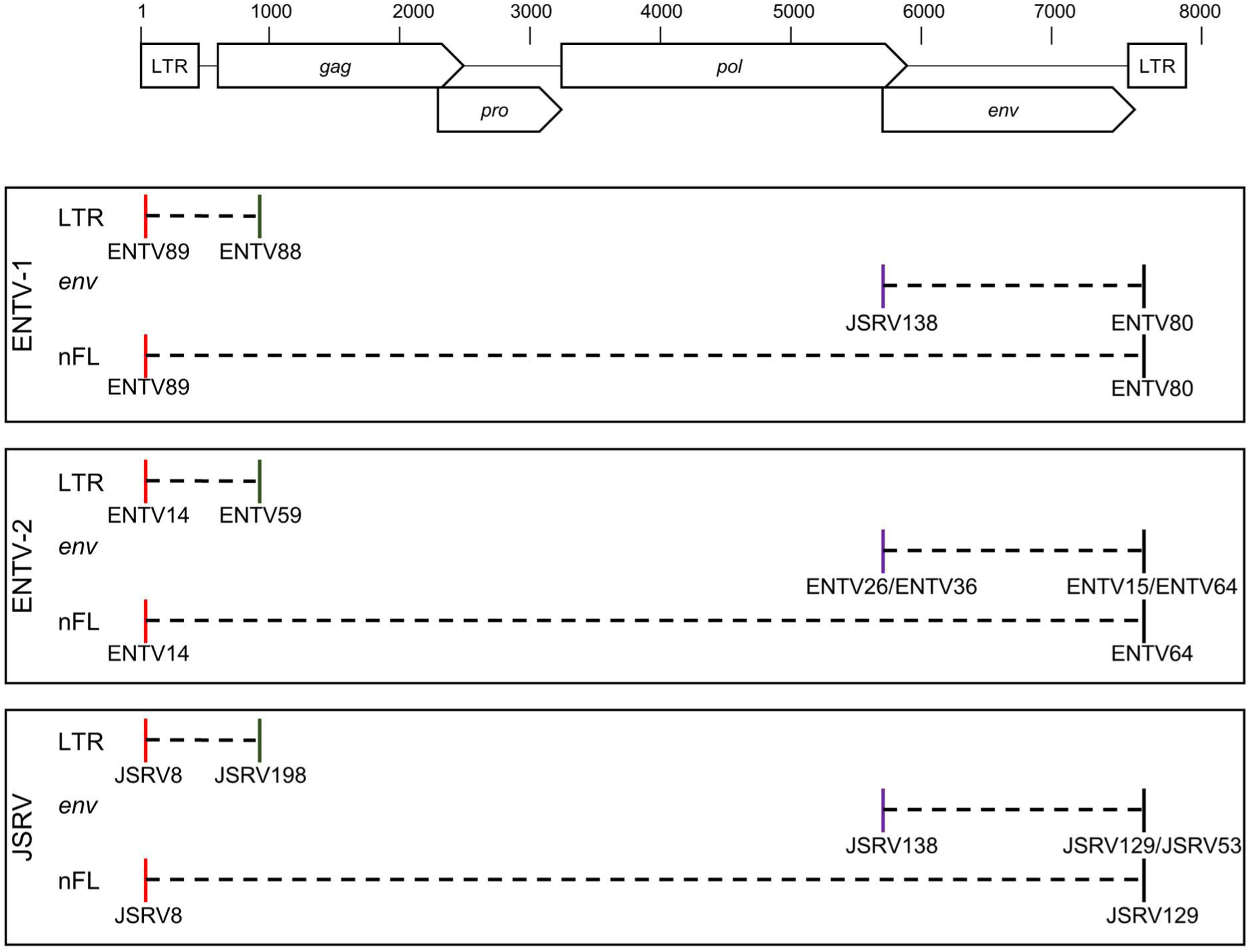
Localization of PCR primers for the amplification of LTR env and near full-length genome. For JSRV and ENTV-2 two different sets of primers were used for the amplification of *env* with the improvement throughout the study of the exo/endo amplification specificity.

**Figure S2:**
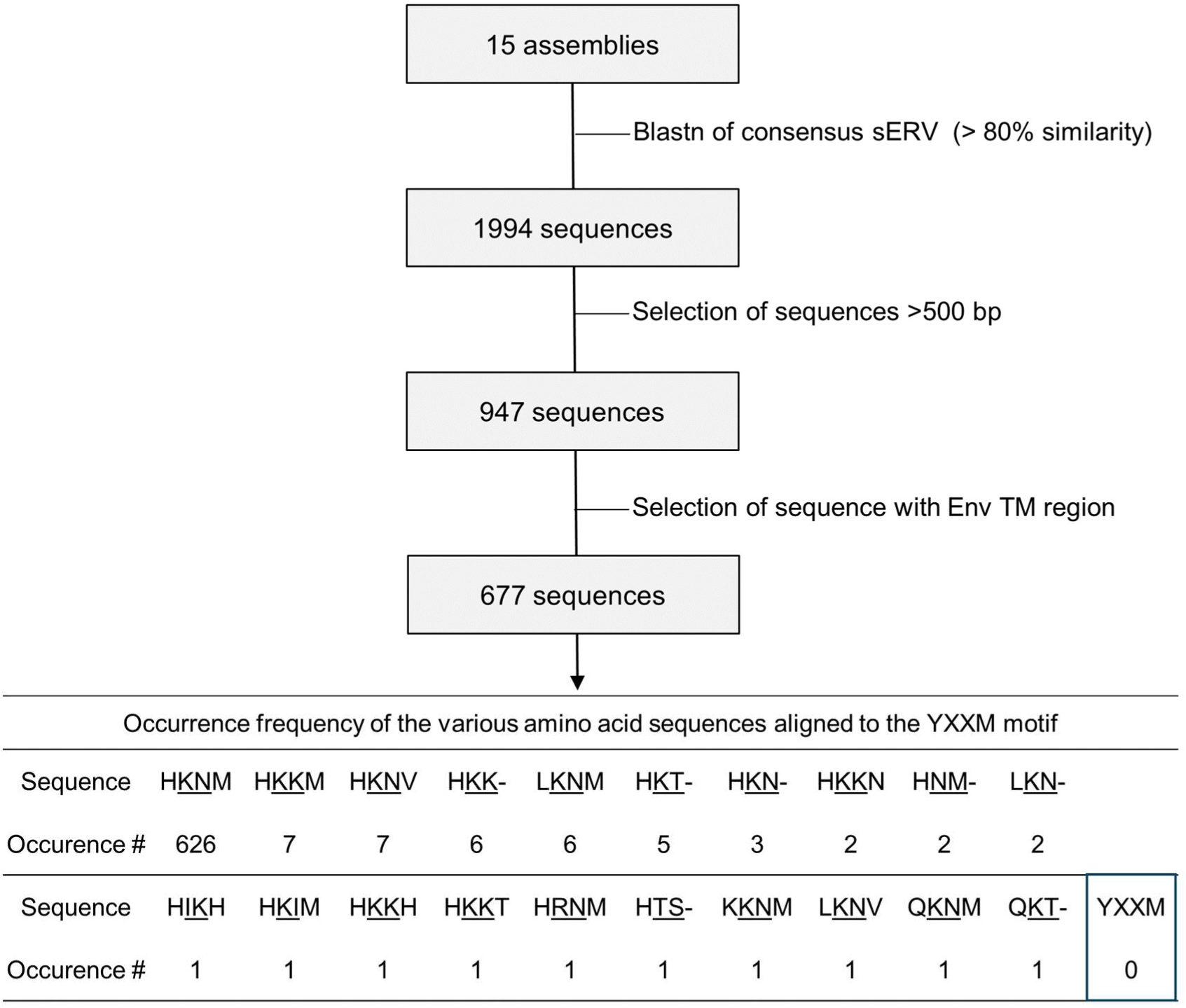
Absence of the YXXM motif in the Env cytoplasmic tail of *caprinae* β-ERVs related to JSRV and ENTVs. sERV and gERV sequences within 15 *caprinae* assemblies (ARS1.2: GCA_001704415.1, ASM1117029: GCA_011170295.1, ASM1914517: GCA_019145175.1, ASM2665220: GCA_026652205.1, CapAeg_1.0: GCA_000978405.1, CAU_F1_maternal_1.0: GCA_023701675.1, CAU_O.aries_1.0: GCA_017524585.1, CVASU_BBG_1.0: GCA_004361675.1, Oar_v4.0: GCA_000298735.2, CAU_Oori_1.0: GCA_014523465.1, ARS-UI_Ramb_v2.0: GCA_016772045.1, Oar_ARS-UKY_Romanov_v1.0: GCA_022244705.1, Saanen_v1: GCA_015443085.1, Oar_ARS-UKY_WhiteDorper_v1.0: GCA_022244695.1, CHIR_2.0: GCA_000317765.2) were identified by BLASTn, using a consensus of sERV from sequences available in GenBank (see figure 1 and Materials and methods). Sequences < 500 bp corresponding to solo LTRs were not included. MAFFT alignment of the different sequences allowed to select only those sequences with a preserved TM region of Env.

**Figure S3:**
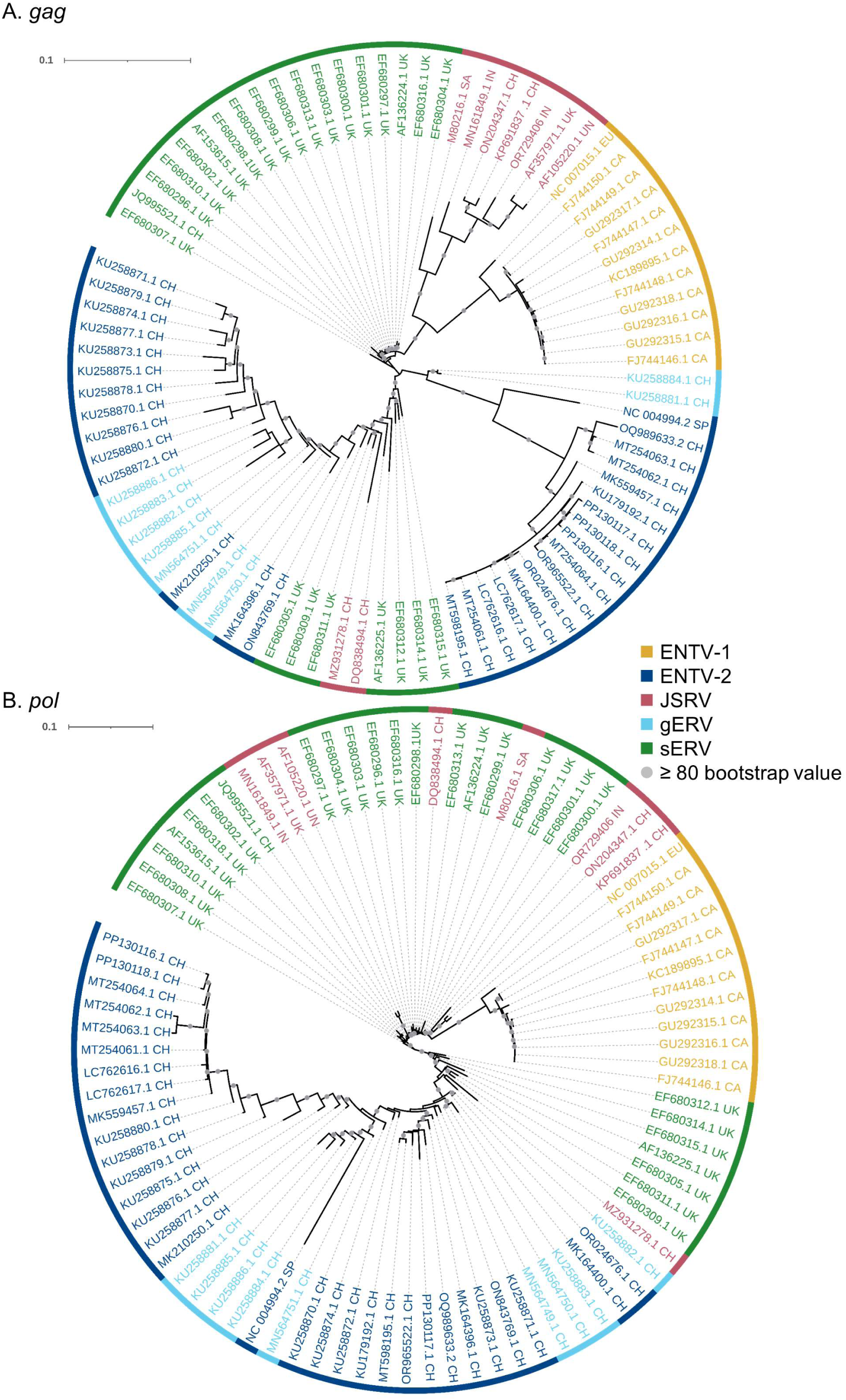
Phylogenetic analysis of sequences of ENTV-1, ENTV-2 and JSRV and their related endogenous sequences from sheep and goats. Maximum likelihood tree with 10,000 ultrafast bootstrap replicated with EF680307.1 (enJSRV10) as root sequence for A) *gag* B) *pol* region. The scale bar indicates the nucleotide substitution per site. CH: China, UK: United Kingdom, CA: Canada, IN: India, SA: South Africa, SP: Spain, EU: Europe, PA: Pakistan, UN: Unknown

**Figure S4:**
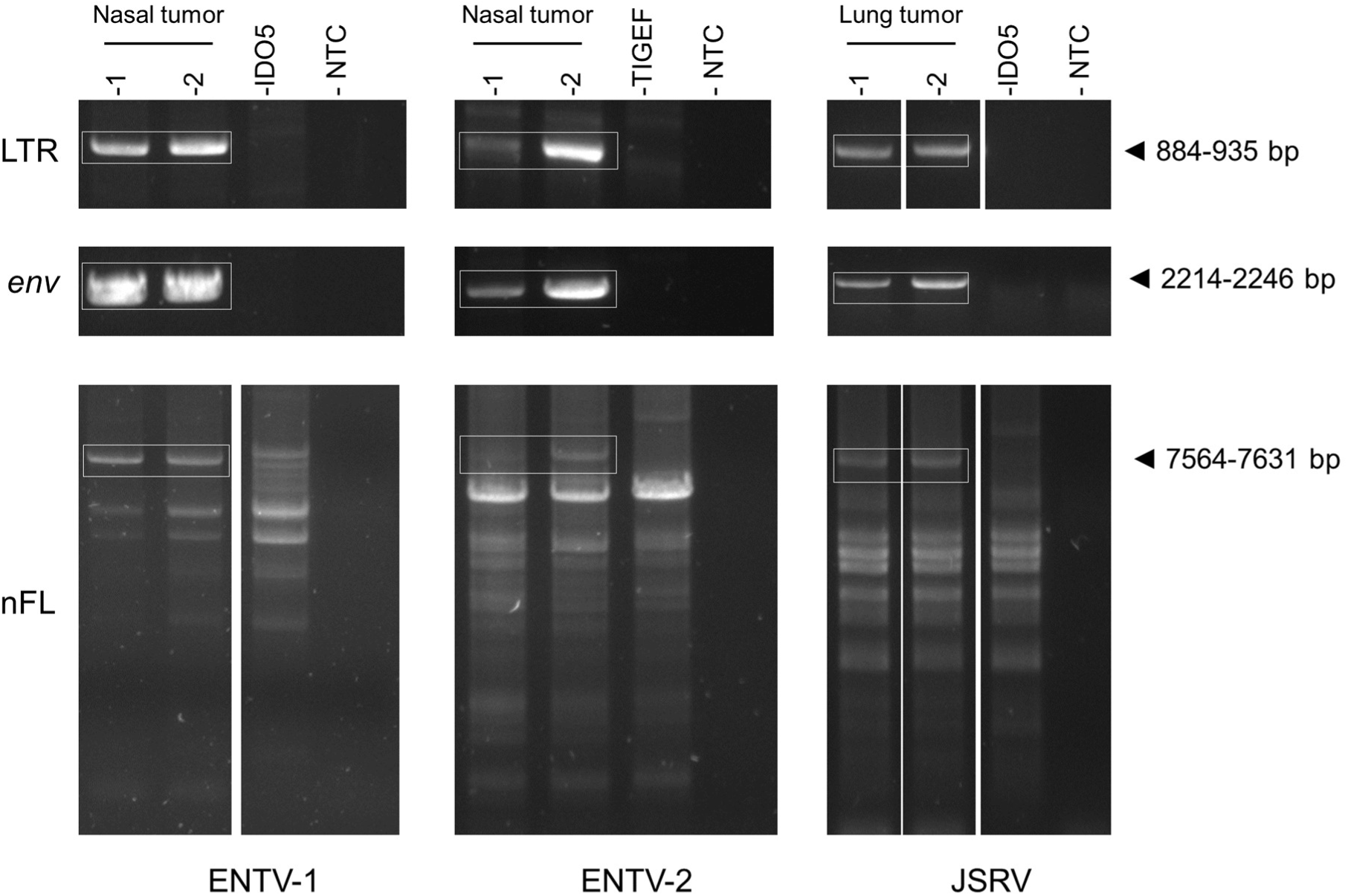
Migration profile of ENTV and JSRV PCR amplification. For each PCR, the example of genomic DNA extracted from two tumors used as a template per virus is shown, in addition to DNA extracted from ovine (IDO5) and caprine (TIGEF) cell lines used as a control for endogenous amplification. nFL = near full-length.

